# Dynamic S-acylation of STIM1 is required for store-operated Ca^2+^ entry

**DOI:** 10.1101/2022.02.15.480522

**Authors:** Goutham Kodkandla, Savannah J West, Qiaochu Wang, Ritika Tewari, Michael X. Zhu, Askar M. Akimzhanov, Darren Boehning

## Abstract

Many cell surface stimuli cause calcium release from endoplasmic reticulum (ER) stores to regulate cellular physiology. Upon ER calcium store depletion, the ER-resident protein STIM1 physically interacts with plasma membrane protein Orai1 to induce calcium release-activated calcium (CRAC) currents that conduct calcium influx from the extracellular milieu. Although the physiological relevance of this process is well established, the mechanism supporting the assembly of these proteins is incompletely understood. Earlier we demonstrated a previously unknown post-translational modification of Orai1 with long chain fatty acids, known as S-acylation. We found that S-acylation of Orai1 is dynamically regulated in a stimulus-dependent manner and essential for its function as a calcium channel. Here we show that STIM1 is also rapidly and transiently S-acylated at cysteine 437 upon ER calcium store depletion. S-acylation of STIM1 is required for the assembly of STIM1 into puncta with Orai1 and full CRAC channel function. Together with the S-acylation of Orai1, our data suggest that stimulus-dependent S-acylation of CRAC channel components Orai1 and STIM1 is a critical mechanism facilitating CRAC channel assembly and function.

## Introduction

Calcium depletion in the endoplasmic reticulum (ER) upon IP_3_-induced calcium release leads to a rapid, highly controlled, and concerted convergence of plasma membrane (PM) calcium channel Orai1 and ER membrane calcium sensing protein STIM1 to form calcium release-activated current (CRAC) channel puncta to promote calcium entry (1). The term puncta is used to define these complexes of Orai1 and STIM1 at the ER-PM junctions (2). Calcium entry through these puncta after store depletion is called store-operated calcium entry (SOCE). The luminal EF hand domain of STIM1 prevents its spontaneous activation by binding to calcium and keeping STIM1 in an inactive conformation (3,4). Dissociation of calcium from the EF hand of STIM1 after store depletion results in a conformational change that exposes its STIM1-ORAI activating region (SOAR) (5). STIM1 then oligomerizes and translocates to PM subdomains where it binds to Orai1 (6,7). Subsequent calcium entry through Orai1 channels is essential for shaping the spatiotemporal aspects of calcium signaling leading to diverse cellular outcomes.

The mechanism that directs Orai1 and STIM1 into CRAC puncta is still incompletely understood. The prevailing hypothesis is a diffusion trap model, which postulates that a STIM1 conformational change leads STIM1 binding to Orai1 in a stochastic manner at ER-PM contact sites to promote SOCE (8,9). STIM1 can bind phosphatidylinositol 4,5-bisphosphate (PIP_2_) via a lysine-rich C-terminal domain, and studies have shown that STIM1 translocates and binds PIP_2_ present specifically in lipid rafts (10). This indicates an active mechanism for targeting STIM1 to lipid rafts, as we and others have shown previously for Orai1 (11,12). Perhaps not surprising, it has also been shown that there is reduced mobility of Orai1 and STIM1 after store depletion at ER-PM contact sites (13,14). However, both proteins are in a state of dynamic equilibrium, and both proteins can “escape” a puncta and join other puncta (8). Put together, the assembly of Orai1 and STIM1 at ER-PM junctions is likely more complicated than a simple diffusion trap model with several factors regulating the assembly and disassembly of the CRAC complex.

S-acylation is a reversible post-translational modification of cysteine residues that is mediated by a specific set of enzymes called palmitoyl acyltransferases (PATs) (15). All known PATs belong to the family of DHHC enzymes owing their name to the aspartate-histidine-histidine-cysteine motif in their catalytic site. S-acylation is known to affect protein stability, trafficking, and recruitment to membrane subdomains (15-17). In contrast to other post-translational lipid modifications like prenylation, myristoylation, or others, S-acylation is reversible and highly labile (15). We have recently shown that Orai1 is dynamically S-acylated after store depletion and during initial stages of T cell activation (11). This was required for Orai1 activation and recruitment into puncta. Here we show that the dynamic S-acylation of STIM1 also mediates SOCE. STIM1 is rapidly and transiently S-acylated after store depletion at cysteine 437, and this is required for recruitment into puncta and SOCE. Our results are consistent with the central role of lipid rafts in mediating the macromolecular assembly of the T-cell receptor (TCR) and other signaling complexes to orchestrate cellular calcium signaling (18).

## Results

### STIM1 is dynamically S-acylated after store depletion at cysteine 437

Previously, we found that S-acylation of Orai1 is increased after stimulation of the T cell receptor (TCR) in Jurkat T cells (11). We tested the hypothesis that STIM1 also undergoes S-acylation after T cell activation. STIM1 has only one cytosolic cysteine residue (C437) that can potentially be a substrate for DHHC enzymes (Fig. 1A). We treated Jurkat T cells with anti-CD3 (OKT3) antibody to stimulate the TCR in these cells. We obtained cell lysates after different time points and selectively enriched the S-acylated proteins using the acyl-resin-assisted capture (acyl-RAC) assay (34). We found that STIM1 is S-acylated after T cell activation which peaked at 5 minutes and then returned to baseline by 15 minutes (Fig. 1B-C). These kinetics are consistent with the activation of the downstream TCR signaling cascade indicated by phosphorylation of PLC-γ1 (Fig. 1D). Next, to show that C437 is the S-acylated residue in STIM1, we transfected HEK293 STIM double knockout (DKO) cells with WT Orai1-Myc and WT or C437S versions of STIM1-Flag. ER calcium store depletion was induced using thapsigargin (TG), and acyl-RAC was performed. Using this approach, we show that WT STIM1, but not the C437S STIM1, is S-acylated after ER calcium store depletion (Fig. 1E). Importantly, the expression levels and ER localization of STIM1 was not compromised by the C437S mutant (Fig. S1). Thus, our data suggests that STIM1 is S-acylated at C437 in a stimulus-dependent manner.

**Figure 1:**
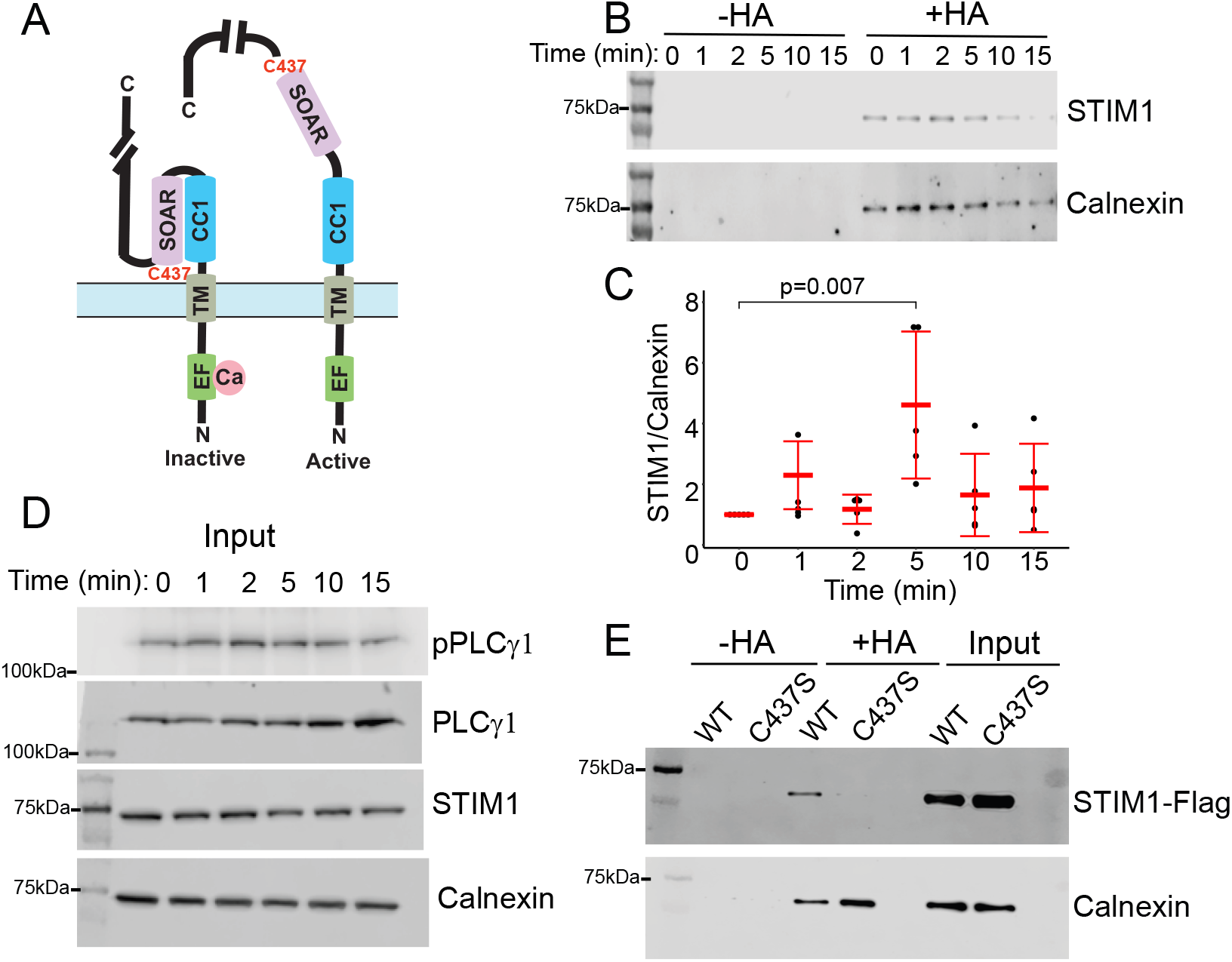
STIM1 is dynamically S-acylated at cysteine 437. A) Schematic representation of the inactive and active (extended) conformation of STIM1. The locations of the EF hand, coiled-coil 1 (CC1) and STIM1 Orai1 activation region (SOAR) of STIM1 are indicated. The location of cysteine 437 is noted at the C-terminus of the SOAR domain. For simplicity, one monomer of the dimeric STIM1 structure is presented. B) Jurkat T cells were treated with anti-CD3 antibody for 0, 1, 2, 5, 10, and 15 minutes and subjected to acyl-RAC. The reaction without hydroxylamine (-HA) serves as a negative control. Calnexin serves both as a positive control for S-acylation and a loading control. Representative of n=5. C) Quantification of fold change of S-acylation of STIM1 normalized to calnexin. Error bars indicate S.D. D) Input blots are shown for levels of pPLCγ1, PLCγ1, STIM1, and calnexin. The pPLCγ1 blot is indicative of the time course of TCR activation. E) HEK STIM double knockout (DKO) cells were transfected with WT or C437S STIM1-FLAG and acyl-RAC was performed after 5 minutes of calcium store depletion with thapsigargin. The blots are representative of four independent experiments.

### S-acylation of STIM1 at cysteine 437 is required for SOCE

To evaluate the effect of S-acylation of STIM1 on CRAC channel function, we monitored whole cell currents in cells co-expressing Orai1-YFP and either WT or the C437S mutant of STIM1-mRFP. Patch pipettes included 50µM IP_3_ to deplete internal stores and activate CRAC currents. As shown in Fig. 2A for current density at −100 mV, cells co-expressing Orai1 and WT STIM1 (grey trace) displayed current developments with three distinct phases: activation (0–45 s), inactivation (50–100 s) and a plateau (100–150 s). In comparison, the activation phase was markedly slowed for cells co-expressing Orai1 and C437S STIM1 (black trace) and the peak current density was also reduced. The current-voltage relationships obtained from voltage ramps at the beginning of whole-cell breaking-in and at the peak of current development showed an inward rectification and reversal potentials at >+40mV (Fig 2B), consistent with the features of CRAC channels. Peak current density and the time to 90% peak at −100mV from multiple cells revealed significant decreases in cells that express C437S STIM1 compared to cells that express WT STIM1 (Fig 2C, D).

**Figure 2:**
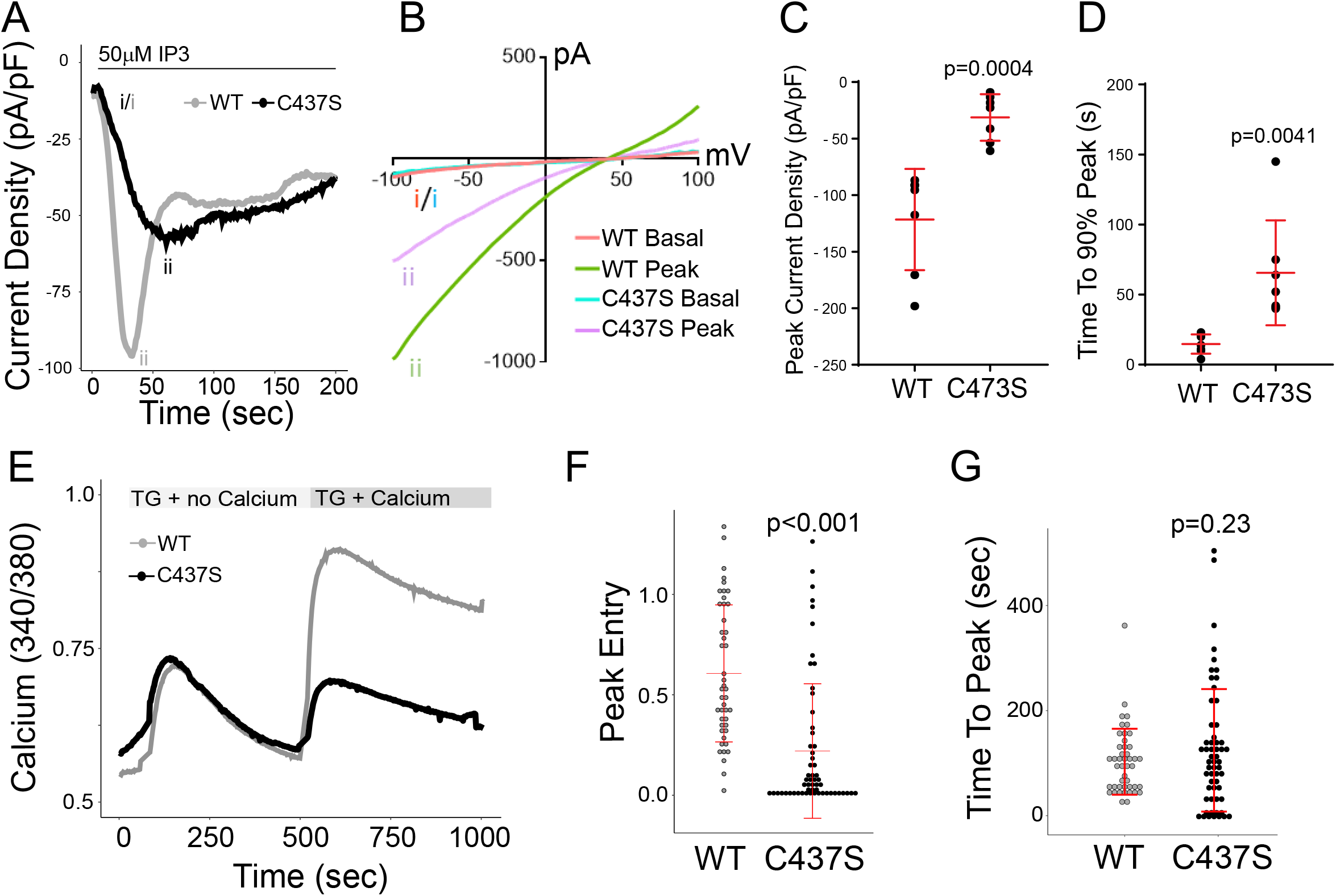
S-acylation of STIM1 facilitates store operated calcium entry. (A) Representative traces of CRAC channel currents in cells expressing WT STIM1-RFP (grey) and the acylation-deficient STIM1 mutant C473S (black). CRAC channel currents (representative of 7 experiments for each condition) were collected at −100 mV immediately upon entering whole-cell configuration. 50 µM IP3 was included in the pipette to activate IP3 receptors to induce ER Ca2+ store depletion. All panels in this figure include co-expression of WT Orai-YFP. (B) Representative current–voltage (I–V) relationships cell expressing WT or C437S STIM1-RFP. Labeling on the left corresponds to the time point that the I–V curve was collected on the trace in A. (C) Quantification of average peak current density of WT and C437S STIM1-RFP expressing cells (n=7 cells for each condition). Error bars indicate S.D. (D) Time to 90% peak current in cells expressing WT and C437S STIM1-RFP. (E) Fura-2 experiments were conducted in STIM double knockout (DKO) HEK293 cells expressing WT or C437S STIM1-RFP. Cells were first imaged in Ca2+-free medium, then treated with 10 µM thapsigargin (TG) in the absence of Ca2+ to induce Ca2+ store depletion. Cells were then incubated with 10 µM TG in the presence of 1 mM Ca2+ to induce Ca2+ entry. Single cell traces are shown as representatives of three individual experiments. There was no difference in basal Ca2+ or peak TG-induced Ca2+ release in the two conditions. (F) Quantification of peak Ca2+ entry in cells expressing WT or C437S STIM1-RFP. (G) Quantification of the time to peak Ca2+ entry in cells expressing WT and C437S STIM1. In E and F, each data point represents a single cell, and the data is pooled from three experiments. Error bars indicate S.D.. The p-values in panels C,D,F, and G are indicated above the C437S data and were calculated with an unpaired two-tailed t-test.

To further evaluate the effect of S-acylation of STIM1 on SOCE, we used Fura-2 imaging in STIM DKO cells. As shown by others, knockout of STIM1 and STIM2 in these cells resulted in a loss of SOCE upon store depletion (Fig. S2; (19)). We rescued STIM1 expression in these cells with either WT or C437S mutant versions of STIM1-mRFP and co-expressed Orai1-YFP. Thapsigargin (TG) in calcium-free buffer was used to induce ER calcium store depletion. Calcium addback was achieved using imaging buffer supplemented with 1mM calcium. Expression of WT STIM1-mRFP rescued SOCE in STIM DKO cells (Fig. 2E). In contrast, cells expressing the C437S version of STIM1-mRFP showed significantly reduced calcium entry (Fig 2E). The peak calcium entry in C437S expressing cells was significantly lower compared to the WT counterpart (Fig. 2F), whereas the time to peak was not affected (Fig. 5G). We conclude that cysteine 437 on STIM1 is required for SOCE.

### S-acylation of STIM1 is required for Orai1/STIM1 assembly

We previously showed that preventing S-acylation of Orai1 results in loss of puncta formation and colocalization with STIM1 upon store depletion (11). We next tested whether S-acylation of STIM1 is required for assembly with Orai1. We co-transfected STIM DKO cells with Orai1-YFP and either WT or C437S versions of STIM1-mRFP and depleted ER stores using 10µM TG. Cells co-transfected with Orai1-YFP and WT STIM1-mRFP showed a rapid and stimulus-dependent colocalization by TIRF imaging (Fig. 3A-C; Fig. S3). In contrast, cells expressing the C437S version of STIM1 showed slower kinetics of colocalization with Orai1 and significantly reduced peak colocalization (Fig. 3A-C; Fig. S3).

**Figure 3:**
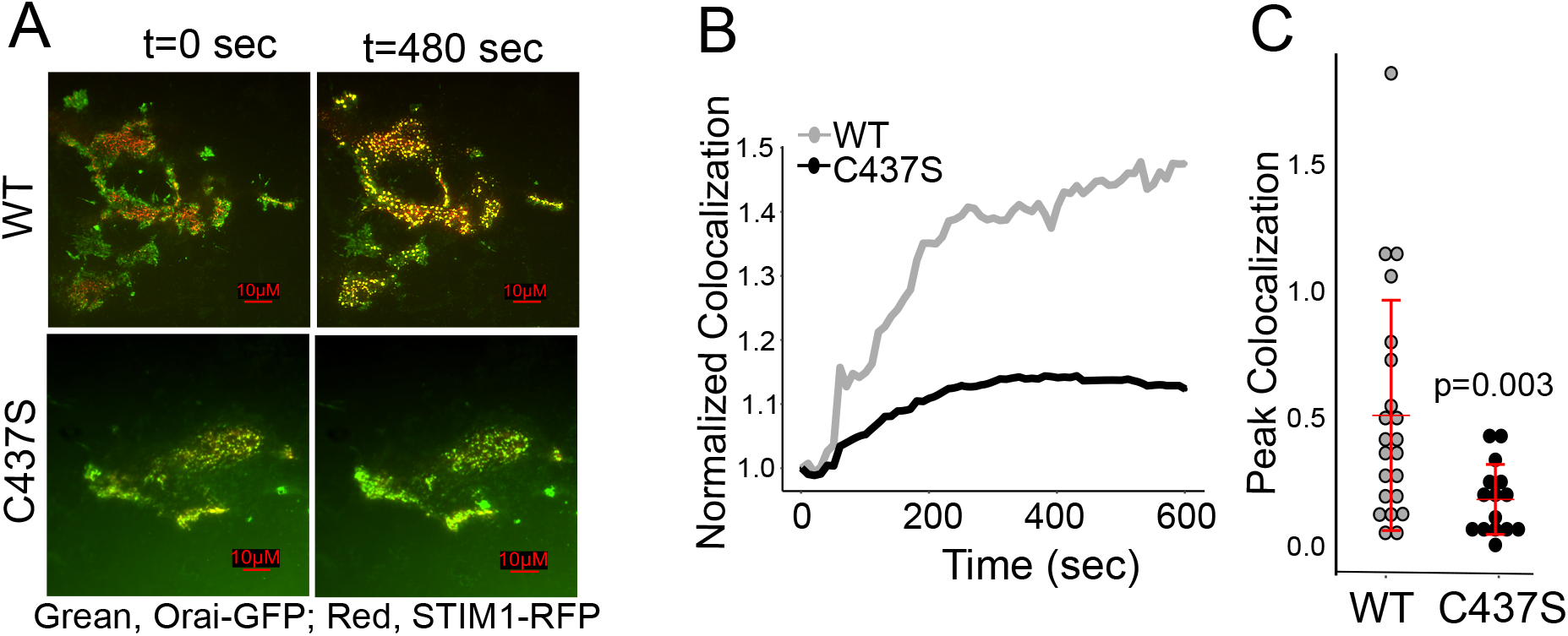
S-acylation of STIM1 is required for colocalization with Orai1. A) Representative TIRF images from six experiments of STIM DKO cells transfected with WT Orai-YFP (green) and either WT (top) or C437S (bottom) STIM1-mRFP (red). Thapsigargin was added after 60 seconds of baseline recording. See also supplementary videos 1 and 2. B) Quantification of WT and C437S STIM1-mRFP colocalization with Orai1-YFP using Pearson’s correlation coefficient over time. Traces represent averages pooled from six separate experiments. The data was normalized to time zero. C) Peak normalized colocalization quantification of WT and C437S STIM1-mRFP and Orai1-YFP. Each data point represents a single cell and the data is pooled from 6 separate experiments. Error bars indicate S.D. The p-value was calculated with an unpaired two-tailed t-test.

### Formation of functional Orai1/STIM1 puncta is compromised by the STIM1 C437S mutant

Orai1 fused to the calcium indicator GCaMP6f allows the direct and local imaging of calcium entry through individual or clusters of Orai1 channels by TIRF imaging (20). We have previously used this approach to investigate how S-acylation of Orai1 affects the formation of active Orai1 puncta (11). To investigate how STIM1 S-acylation affects the Orai1 channel activation and puncta formation, we co-transfected STIM DKO cells with WT Orai1-GCaMP6f and either the WT or C437S version of STIM1-mRFP. The addition of 10µM TG in calcium free media led to puncta formation in WT STIM1 expressing cells, with much fewer puncta in C437S expressing cells (Fig. 4A-C; Fig. S4). Thapsigargin treatment in calcium free media resulted in no increase in Orai1-GCaMP6f fluorescence in either WT or C437S STIM1 expressing cells, consistent with the specificity of this protein in measuring calcium entry through the mouth of the channel (Fig. 4A-B; Fig. S4). Calcium addback caused a significant increase in Orai1-GCaMP6f fluorescence in cells expressing WT STIM1, indicative of calcium entry through Orai1 channels. In contrast, cells expressing C437S STIM1 did not show a significant increase in fluorescence after calcium addback, but only recovered to the level comparable to that of before extracellular calcium removal (Fig 4A-B,D; Fig. S4). We conclude that the STIM1-C437S mutant has defects in both the recruitment to Orai1 puncta and gating the channel.

**Figure 4:**
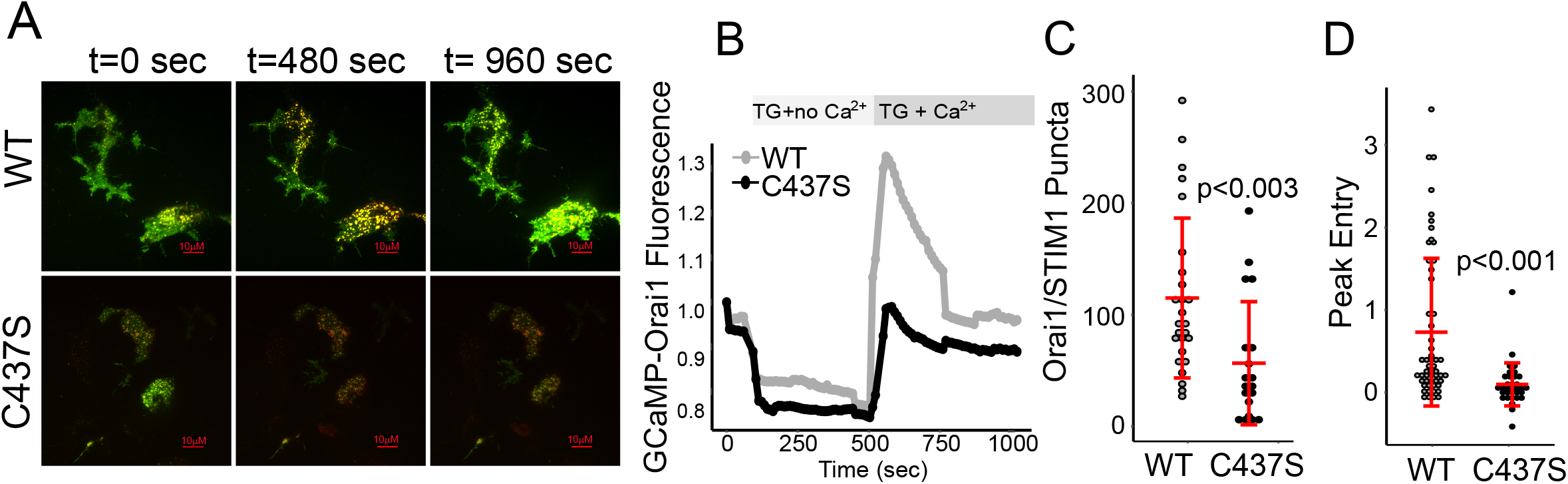
STIM1 S-acylation is required for the recruitment of active Orai1 to puncta. A) Representative TIRF images from six experiments of STIM DKO cells transfected with WT Orai-GCaMP6f (green) and either WT (top) or C437S (bottom) STIM1-mRFP (red). Cells were first imaged in Ca2+-free medium, then treated with 10 µM thapsigargin (TG) in the absence of Ca2+ to induce Ca2+ store depletion. Cells were then incubated with 10 µM TG in the presence of 1 mM Ca2+ to induce Ca2+ entry. See supplementary videos 3 and 4. B) Normalized Orai-GCaMP6f fluorescence of cells transfected with WT and C437S STIM1-mRFP over time. Traces represent averages pooled from six separate experiments. C) Quantification of the number of Orai1–GCaMP6f/STIM1 puncta which appear after Ca2+ add-back. D) Peak normalized fluorescence of Orai1-GCaMP6f in cells expressing WT and C437S STIM1-mRFP. Each data point in Panels C,D represents a single cell and the data is pooled from 6 separate experiments. Error bars indicate S.D. The p-values were calculated using an unpaired two-tailed t-test.

## Discussion

Store-operated calcium entry is an important mechanism for calcium refilling after ER calcium store depletion and contributes significantly to shaping the spatiotemporal aspects of calcium signaling (21,22). The mechanisms behind how the CRAC channel components are translocated to the ER-PM junctions to form puncta are not fully understood. Previously, our group demonstrated that Orai1 is dynamically S-acylated after store depletion and a cysteine mutant version of this protein that cannot undergo S-acylation effects CRAC channel assembly and SOCE (11), findings that have also been validated by others (12). Here we show that S-acylation of STIM1 also plays a crucial role in CRAC puncta formation and that it contributes to SOCE.

The S-acylation of Orai1 in the PM targets the channel to lipid rafts where it forms CRAC channels with STIM1 to promote calcium entry (11,12). We hypothesize S-acylation of STIM1 regulates SOCE in two possible ways. One model postulates that STIM1 is S-acylated by an ER-localized DHHC enzyme to promote and/or stabilize the extended conformation of STIM1 (Fig. 5). Another possible model hypothesizes the S-acylation of the C-terminus of STIM1 in the extended conformation by a PM-localized DHHC enzyme. In this model, S-acylation would function not only to anchor the SOAR/CAD domain of STIM1 in the PM but also promote localization to lipid rafts to promote binding to raft-localized Orai1 (Fig. 5). Importantly, this model proposes that both the assembly and disassembly of CRAC channels is an enzymatically regulated process. If this model is correct, a significant revision to the diffusion-trap model of CRAC channel assembly would be required.

**Figure 5:**
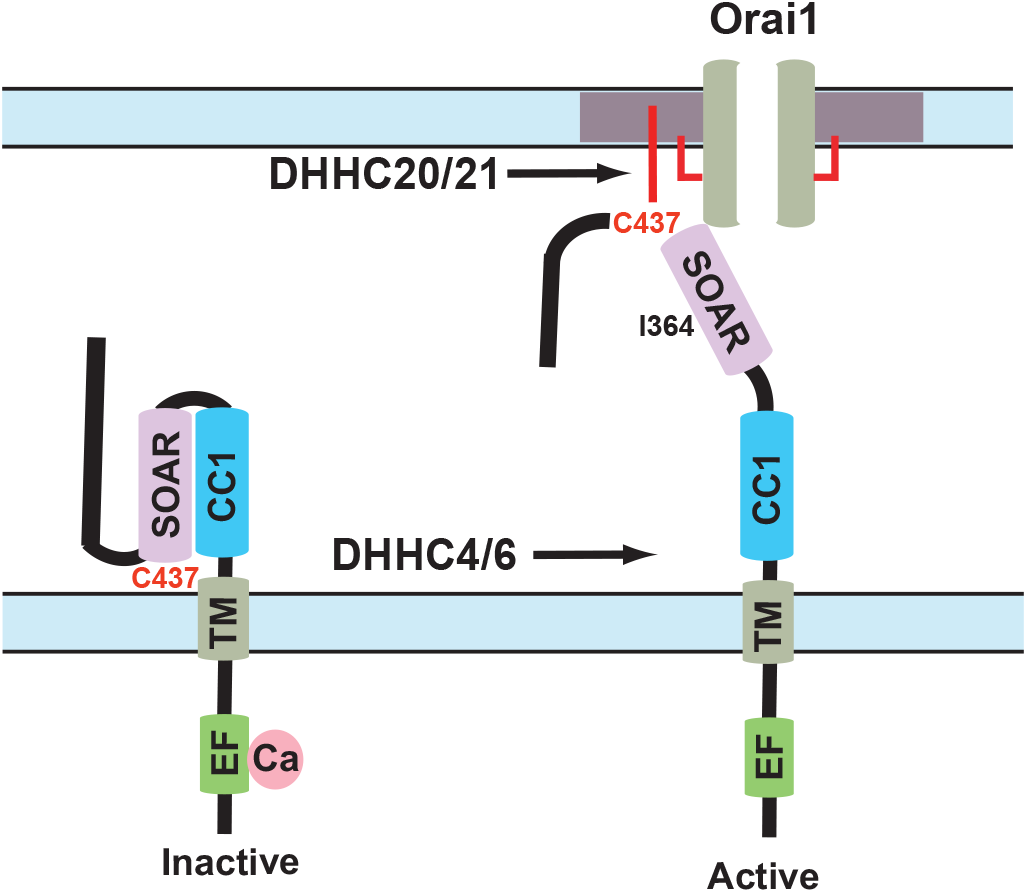
S-acylation of STIM1 C437 and CRAC channel function. Two potential models for the regulation of STIM1 by S-acylation at C437. In the first model, ER-localized DHHC4 or DHHC6 S-acylate C437 facilitating and/or stabilizing the extended conformation of STIM1. In the second model, S-acylation of C437 by the plasma membrane-localized DHHC20 or DHHC21 both stabilizes the extended conformation and promotes recruitment to lipid rafts where it can bind and gate S-acylated Orai1. The cholesterol binding residue isoleucine 364 is also highlighted. For clarity, the C-terminal PIP2 binding polybasic domain is not shown.

Active STIM1 adopts an extended conformation upon store depletion. This fully extended conformation facilitates binding of the polybasic domain of STIM1 to the PIP_2_-rich domains specifically in PM lipid rafts (10). In addition, STIM1 has a cholesterol binding domain in the SOAR region, of which isoleucine 364 plays a critical role (23). Thus, in addition to S-acylation, there are multiple structural features in the C-terminus of STIM1 which promotes binding and segregation to lipid rafts. This redundancy in function is a likely explanation for the partial loss of SOCE we describe here with the C437S mutant STIM1. Interestingly, it has been shown that STIM1 encompassing only residues 1-442 still forms puncta in Orai1 triple knockout cells, whereas STIM1 encompassing residues 1-342 does not (24). These results indicate that Orai1 and the polybasic domain of STIM1 are dispensable for puncta formation, however the SOAR domain (containing C437 and I364 residues) is absolutely required.

It is possible that the same plasma membrane DHHC enzyme mediates the S-acylation of both Orai1 and STIM1. This hypothesis is supported by our findings that both Orai1 and STIM1 reach peak S-acylation levels at the same time point after T cell stimulation (Fig. 1B-C; (11)). Similarly, we have observed in TIRF time lapse imaging experiments that the peak colocalization of Orai1 and STIM1 happens approximately at the same time after store depletion (Fig. 3B; (11)). It has been shown that DHHC20 S-acylates Orai1 (12). We have shown that many TCR pathway components are S-acylated by the PM-localized DHHC21 (25-28). Future work will evaluate if Orai1 can also be S-acylated by DHHC21 and determine the DHHC enzyme(s) responsible for STIM1 S-acylation. Of note, we have shown dramatic defects in T cell function in DHHC21 mutant mice *in vitro* and *in vivo* which may be consistent with alterations in SOCE (25).

Another important observation from S-acylation of STIM1 and Orai1 is the importance of immunological synapse formation in the TCR-mediated immune response (29-31). S-acylation of Orai1 is important for segregation of the channel to the immunological synapse (12). In our previous studies, we have shown that several TCR components are rapidly and transiently S-acylated during T cell activation to regulate immune responses (25-27,32). T cell signaling proteins such as Lck and ZAP70 are recruited into the immunological synapse, and we have shown that S-acylation is an important mechanism behind this recruitment and activation (26,28). We can now include the Orai1/STIM1 complex in the dynamically S-acylated proteome of T cells. As such, CRAC components are co-localized to the immunological synapse with TCR signaling proteins following cellular stimulation to promote sustained calcium entry that is required for immune responses (1,31). Our results also indicate that the S-acylation enzymatic machinery may be a viable therapeutic target for diseases of immune cell function.

## Material and Methods

### Cells, antibodies, and constructs

HEK293 STIM DKO cells were a kind gift of Dr. Mohamed Trebak (19) (University of Pittsburgh, Pittsburgh, PA, USA) and were maintained in Dulbecco’s modified eagle medium (DMEM) supplemented with 10% fetal bovine serum, 1% L-glutamine, and 1% penicillin-streptomycin. Jurkat T cells were obtained from ATCC and cultured in RPMI media supplemented with 10% fetal bovine serum, 1% L-glutamine, 1% penicillin-streptomycin. Cells were plated on polystyrene tissue culture dishes or 6 well plates for experiments. For some experiments, cells were plated on poly-L-lysine coated coverslips for imaging. Cells were transfected with Lipofectamine 3000 (Invitrogen) according to manufacturer’s protocol with 0.75µg Orai1 plasmid and/or 2.25µg STIM1 plasmid per 35mm well. Cells were maintained at 37ºC and 5% CO2. For immunoblotting, the antibodies were purchased from commercially available sources: STIM1 (#4961S), calnexin (#2679), phospholipase Cγ1 (PLCγ1) (#2822S), phospho-PLCγ1 (Tyr783) (#2821S), anti-rabbit IgG, HRP-linked secondary antibody (#7074S), anti-mouse IgG, HRP-linked secondary antibody (#7076P2) were from Cell Signaling Technology; anti-CD3 human antibody (14-0037-82) was from eBiosciences. Orai1-YFP was a generous gift from Anjana Rao (Addgene plasmid #19756). Orai1-GCamp6f was a plasmid deposited in Addgene by Michael Cahalan (Addgene #73564; (20)). STIM1-mRFP was a generous gift from David Holowka and Barbara Baird (Cornell University, Ithaca, NY, USA). The C437S mutant version of STIM1-mRFP and STIM1-Flag were prepared using q5 site-directed mutagenesis kit from New England Biolabs. The following primers for mutagenesis were designed using the NEB tool 5’-GAGATCAACCTTGCTAAGCAGG-3’ and 5’-GAAACACACTCTTTGGCACCTT-3’. The mutation was confirmed using Sanger sequencing (Eton Biosciences). Thiol-sepharose beads used for acyl-RAC were obtained from Nanocs Inc and activated according to manufacturer’s protocol before continuing with acyl-RAC protocol listed below. All other chemicals and reagents were purchased from Sigma-Aldrich or VWR.

### Electrophysiology

We performed whole-cell recording using HEK293 cells co-expressing Orai1-YFP with WT or C437S STIM1-mRFP as described previously(11). Patch-clamping was conducted with an EPC-10 amplifier controlled by PatchMaster software (HEKA) within 24 hours after transfection. The resistance of fire polished pipettes was between 5 megohm to 8 megohm. The pipette solution contained 120mM cesium glutamate, 20mM cesium, BAPTA [1,2-bis(2-aminophenoxy)ethane-N,N,N’,N’-tetraacetic acid], 3mM MgCl_2_, 0.05mM IP_3_ (K^+^-salt), and 10mM HEPES (pH 7.2 with CsOH). The bath solution contained 120mM NaCl, 10mM CaCl_2_, 2 mMMgCl_2_, 10mM TEA-HCl (tetraethylammonium chloride), 10mM glucose, and 10mM HEPES (pH 7.2 with NaOH). Cells were held at the holding potential of 0mV and currents recorded at a sampling frequency of 10kHz with 1kHz filtering. Currents were elicited by voltage ramps from −100 to +100mV in 100mS, repeated every second. To capture current development immediately after IP_3_ dialysis, the recording began before the establishment of the whole-cell configuration and continued throughout the breaking-in and afterwards. Patches with seal resistance up to 3 gigohm to 5 gigohm were broken by applying negative pressure and only cells that maintained gigohm resistance after the breaking-in were used for analysis. All voltages were corrected for a liquid junction potential of −10 mV.

### Fura-2 Imaging

Calcium imaging on HEK293 cells was performed as previously described (33). Briefly, two days before imaging, approximately 300,000 HEK293 STIM DKO cells were seeded on coverslips in a 6 well culture plate. After 24h, these cells were transfected with WT Orai1-YFP and either WT or C437S STIM1-mRFP plasmids. All calcium imaging was performed in 1% BSA, 107mM NaCl, 20mM HEPES, 2.5mM MgCl_2_, 7.5mM KCl, and 11.5mM glucose with or without the presence of 1mM CaCl_2_. Time-lapse images were obtained every two seconds for 16 minutes using a 40X oil immersion objective lens. Cells were excited using 340nm and 380nm wavelengths alternatively every two seconds and emission was collected at 525nm. The first minute of recording was used as baseline. After 1 min, 10µM thapsigargin (TG) in calcium free imaging solution was added to induce ER calcium store depletion. This was continued for 7 minutes and imaging solution with calcium and thapsigargin was added back to induce calcium entry. The ratio of fluorescence at 340nm to 380nm was used to quantify cytosolic calcium. We excluded the cells which did not respond to thapsigargin in our analysis. For calculating Peak Entry, we subtracted the maximum fluorescence ratio (R_max_) after calcium addback from the fluorescence ratio at time zero (R_0_) and normalized to R_0_.

### Confocal imaging

HEK293 STIM DKO cells were plated on coverslips in a 6 well plate at a density of 350,000 cells per coverslip. Next day, they were transfected with WT or C437S version of STIM1-mRFP using Lipofectamine 3000 per manufacturer’s protocol. The next day, cells were imaged using Nikon Ti2 confocal microscope with a plan apo lambda 60x oil objective at 1.5 magnification factor. Exposure parameters were identical between the WT and STIM1 expressing cells.

### Acyl-resin-assisted capture (acyl-RAC) assay

We performed the acyl-RAC assay as described by our group elsewhere (34). Cells were transfected with WT Orai1-Myc and either WT or C437S version of STIM1-Flag plasmids. For Jurkat T cells, 10 million cells per treatment were used for endogenous STIM1 S-acylation experiments and were treated with 5µg/ml anti-CD3 (OKT3) antibody. For STIM1 exogenous expression paradigms, we used 10µM thapsigargin for 5 minutes to induce ER calcium store depletion. Cell lysates were collected after the specific time points using 1% dodecyl ß-D-maltoside in PBS, supplemented with cOmplete protease inhibitor cocktail (Roche), acyl protein thioesterase inhibitor ML211, and serine protease inhibitor PMSF (10mM). The lysates were cleared at 20,000 xg for 30 minutes at 4ºC by centrifugation. Approximately 500 µg of cell lysates were used for the assay. The lysates were precipitated using 2:1 methanol and chloroform, and the resulting protein pellets were incubated with 0.2% methylmethanethiosulfonate (MMTS) for 15 minutes at 42ºC. MMTS was removed using three rounds of methanol chloroform precipitation. After the last round of precipitation, the pellets were dissolved in 2SHB buffer (2% SDS, 5mM EDTA, 100mM HEPES, pH 7.4). 20µl of lysate was saved for input. The lysates were incubated with 400mM hydroxylamine (HA) to cleave the thioester bonds and incubated with thiolsepharose resin overnight at 4ºC. For the minus HA controls, lysates were incubated with 400mM sodium chloride in place of HA. Next day, the samples were washed four times using 1% SDS solution in Buffer A (5mM EDTA, 100mM HEPES, pH 7.4) and eluted using 10mM dithiothrietol (DTT) in SDS buffer (1% SDS, 50mM Tris-HCl, 10% glycerol, 1% Bromophenol Blue). The samples were resolved on 10% SDS-PAGE gels and analyzed using western blotting.

### Total internal reflection imaging (TIRF)

HEK293 STIM DKO cells were seeded on coverslips treated with poly-L-lysine two days before imaging at 300,000 cells per coverslip. Next day, these cells were transfected with WT Orai1-YFP or either WT or C437S STIM1-mRFP. For TIRF imaging, we used a Nikon Eclipse Ti microscope equipped with TIRF illumination system and a 60X oil objective. The cells were mounted on the microscope and alternatively excited using 488nm and 561nm lasers every ten seconds. The images were acquired for 16 minutes using a Photometrics Prime 95B camera. The first minute of the recording was used as baseline. Thapsigargin in calcium free imaging solution was added after the first minute to observe Orai1 and STIM1 PM targeting and recruitment to CRAC puncta. For the experiments in Fig. 4, we used Orai1-GCaMP6f with WT or C437S version of STIM1-mRFP to evaluate the effect of STIM1 S-acylation on Orai1 channel activity. We altered the protocol to evaluate calcium entry and imaged every 5 seconds. Imaging was done for 16 minutes and the first minute was used as baseline. After 1 minute, thapsigargin in calcium free imaging solution was added to observe STIM1 colocalization with Orai1. After 8 minutes, the solution was replaced with imaging solution with calcium to observe calcium entry. We excluded all cells with aggregates under resting conditions from our analyses. We used Nikon NIS Elements software for fluorescence data analysis. We drew regions of interest (ROI) around cells to calculate relative fluorescence units (RFU) at any point during the timelapse. We normalized the RFU of specific cell to itself by dividing a specific timepoint fluorescence (Cn) value to time zero (C0). We use these normalized fluorescence values to plot Orai1 channel fluorescence over the time course of thapsigargin addition and calcium addback. To calculate the peak entry, we obtained the maximum fluorescence value (Cmax) after calcium addback and subtracted it from the fluorescence value at time zero (C0) and normalized the difference by dividing it to time zero (C0) (Cmax-C0/C0). To calculate the colocalization between Orai1 and STIM1, we used colocalization analysis on Nikon NIS Elements software. The software uses Pearson’s correlation coefficient as an estimate for colocalization between two fluorophores. We drew ROIs around cells of interest and obtained correlation coefficients for the time series of all cells. We followed the similar method of normalized correlation by dividing correlation coefficient at a given time point (Rn) by time zero (R0). We used these values to plot the colocalization curve. To calculate peak colocalization, we used the similar method to that of peak entry, we subtracted the peak colocalization after thapsigargin addition (Rmax) from time zero (R0) and divided the difference to time zero (R_0_) (Rmax-R0/R0).

---

**Figure S1:**
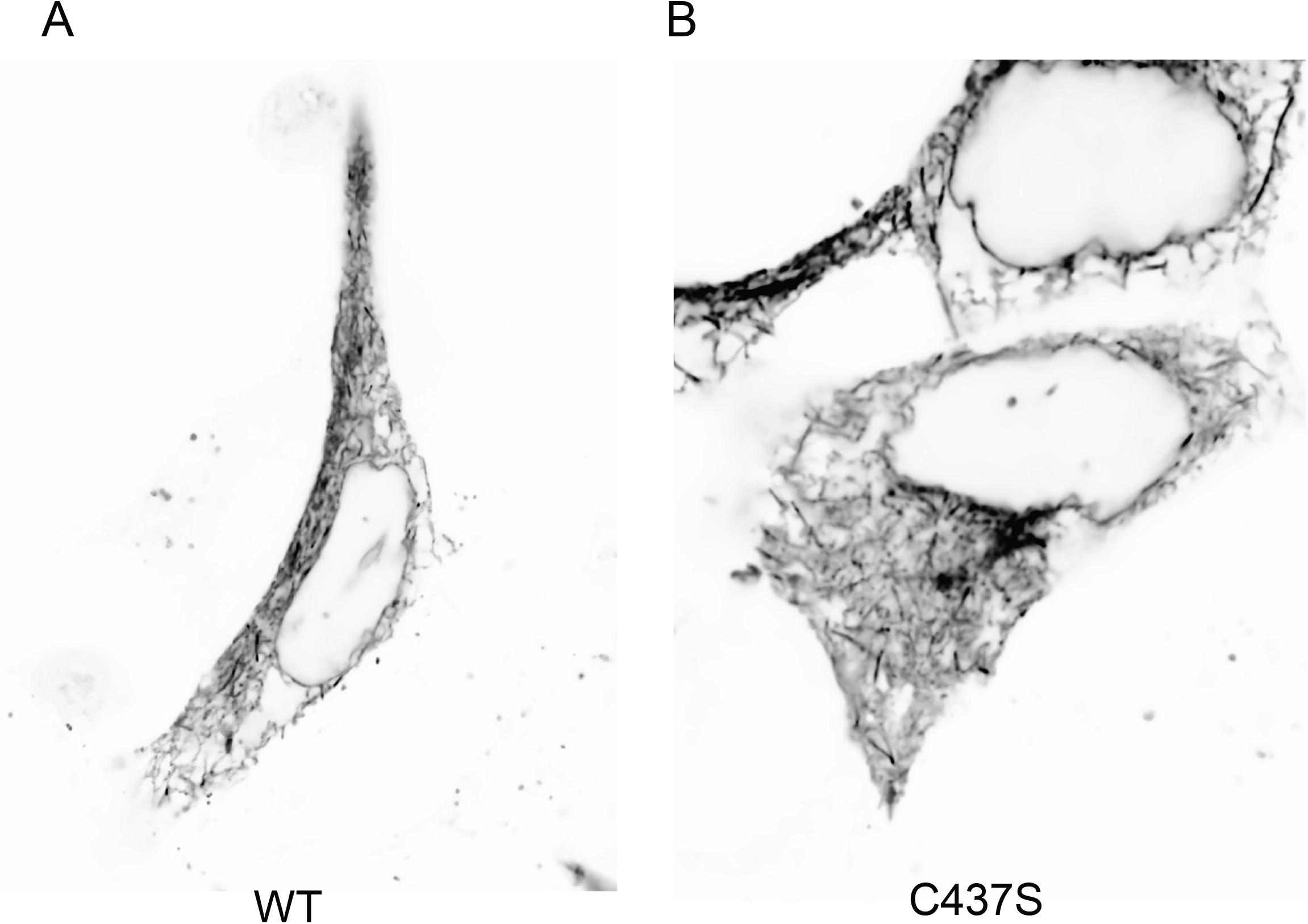
ER localization of WT and C437S STIM1. HEK293 STIM DKO cells were transfected with either WT or C437S STIM1-RFP and imaged by confocal microscopy. An inverted LUT was applied. Images were obtained with a 60X objective with 1.5X intermediate magnification.

**Figure S2:**
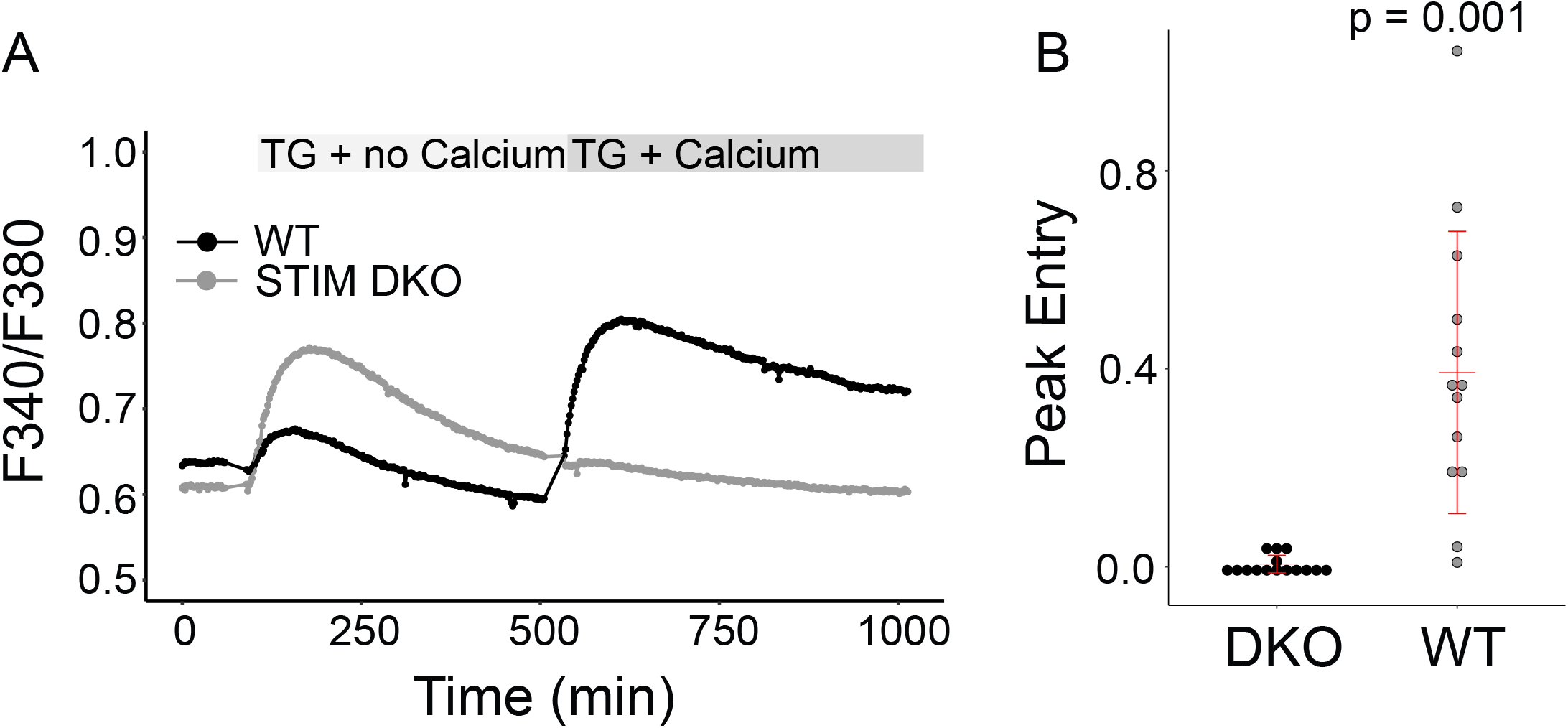
STIM1 DKO cells do not have SOCE. A) Fura-2 imaging of WT and STIM DKO HEK293 cells. Cells were first imaged in Ca2+-free medium, then treated with 10 µM thapsigargin (TG) in the absence of Ca2+ to induce Ca2+ store depletion. Cells were then incubated with 10 µM TG in the presence of 1 mM Ca2+ to induce Ca2+ entry. B) Quantified peak entry.

**Figure S3:**
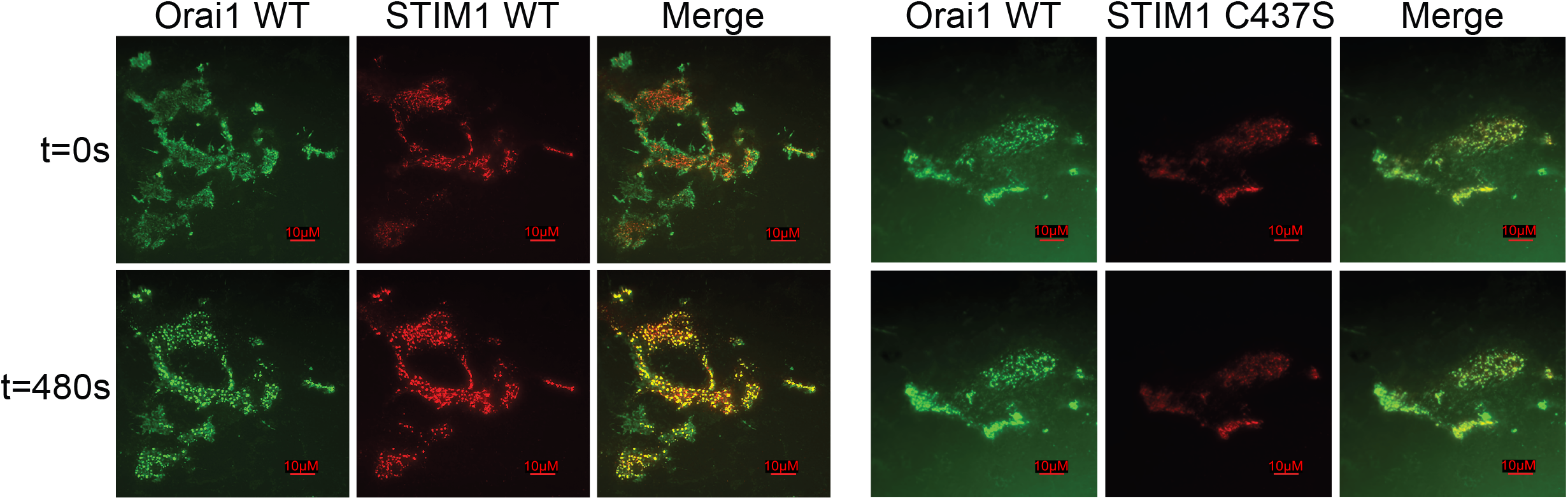
Individual channels from Figure 3. Shown are individual Orai1-YFP, STIM1-mRFP WT, and STIM1-mRFP C437S channels obtained by TIRF microscopy as presented in Figure 3. Time (in seconds) is shown to the left of the panels. Thapsigargin (10µM) was added at 60 seconds. A 10µm scale bar is shown.

**Figure S4:**
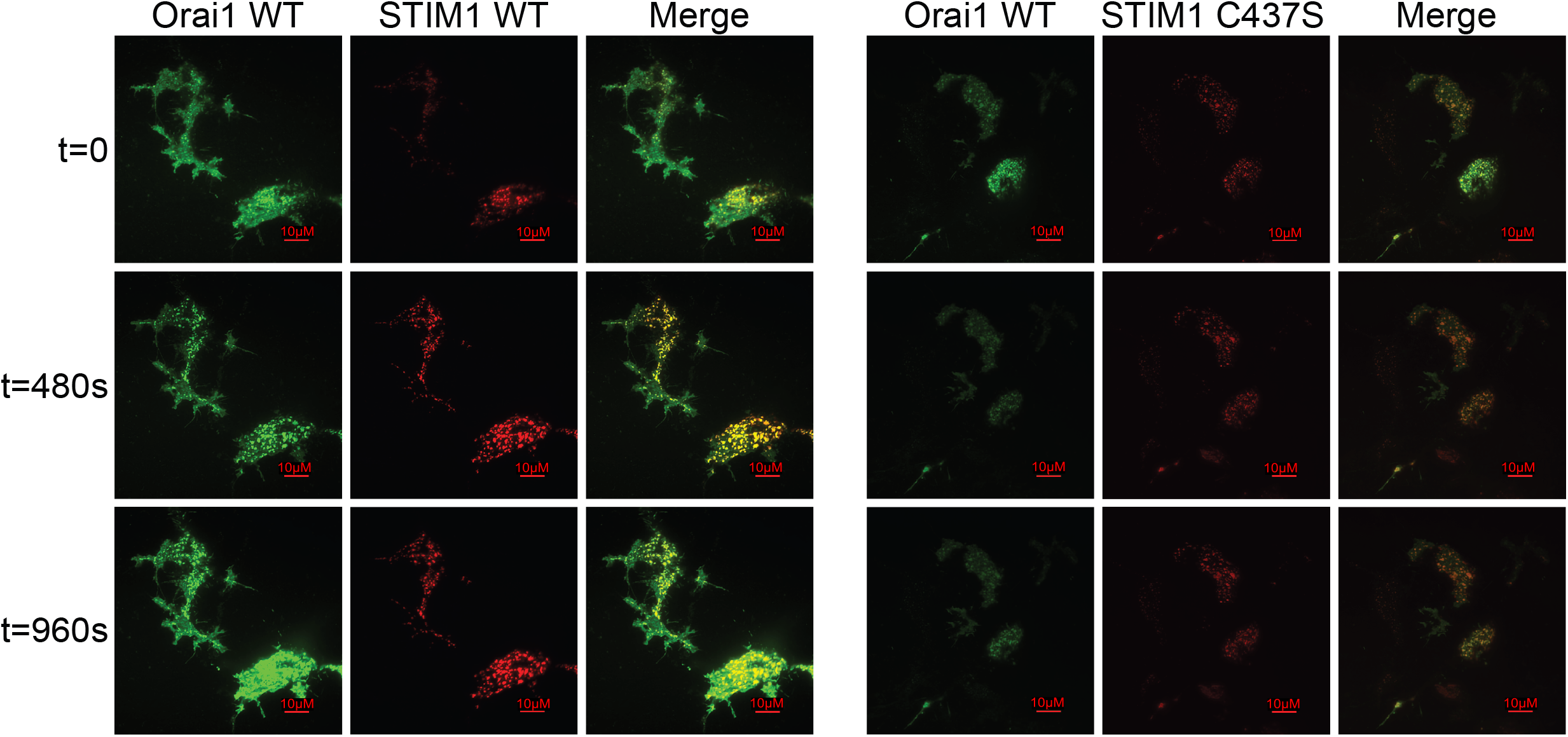
Individual channels from Figure 4. Shown are individual Orai1-GCaMP6f, STIM1-mRFP WT, and STIM1-mRFP C437S channels obtained by TIRF microscopy as presented in Figure 4. Time (in seconds) is shown to the left of the panels. Thapsigargin (10µM) in calcium-free media was added at 60 seconds. Calcium add-back with thapsigargin was at 500 seconds. A 10µm scale bar is shown.

## Notes

### Competing Interest Statement

The authors have declared no competing interest.

### Summary of Updates

Updated author name Updated fig2 labels

## References

1. Lioudyno, M. I., Kozak, J. A., Penna, A., Safrina, O., Zhang, S. L., Sen, D., Roos, J., Stauderman, K. A., and Cahalan, M. D. (2008) Orai1 and STIM1 move to the immunological synapse and are up-regulated during T cell activation. Proc Natl Acad Sci U S A 105, 2011–2016

2. Barr, V. A., Bernot, K. M., Srikanth, S., Gwack, Y., Balagopalan, L., Regan, C. K., Helman, D. J., Sommers, C. L., Oh-Hora, M., Rao, A., and Samelson, L. E. (2008) Dynamic movement of the calcium sensor STIM1 and the calcium channel Orai1 in activated T-cells: puncta and distal caps. Mol Biol Cell 19, 2802–2817

3. Gudlur, A., Zeraik, A. E., Hirve, N., Rajanikanth, V., Bobkov, A. A., Ma, G., Zheng, S., Wang, Y., Zhou, Y., Komives, E. A., and Hogan, P. G. (2018) Calcium sensing by the STIM1 ER-luminal domain. Nat Commun 9, 4536

4. Liou, J., Kim, M. L., Heo, W. D., Jones, J. T., Myers, J. W., Ferrell, J. E., and Meyer, T. (2005) STIM is a Ca2+ sensor essential for Ca2+-store-depletion-triggered Ca2+ influx. Curr Biol 15, 1235–1241

5. Hirve, N., Rajanikanth, V., Hogan, P. G., and Gudlur, A. (2018) Coiled-Coil Formation Conveys a STIM1 Signal from ER Lumen to Cytoplasm. Cell Rep 22, 72–83

6. Kim, J. Y., and Muallem, S. (2011) Unlocking SOAR releases STIM. EMBO J 30, 1673–1675

7. van Dorp, S., Qiu, R., Choi, U. B., Wu, M. M., Yen, M., Kirmiz, M., Brunger, A. T., and Lewis, R. S. (2021) Conformational dynamics of auto-inhibition in the ER calcium sensor STIM1. Elife 10

8. Wu, M. M., Covington, E. D., and Lewis, R. S. (2014) Single-molecule analysis of diffusion and trapping of STIM1 and Orai1 at endoplasmic reticulum-plasma membrane junctions. Mol Biol Cell 25, 3672–3685

9. Hoover, P. J., and Lewis, R. S. (2011) Stoichiometric requirements for trapping and gating of Ca2+ release-activated Ca2+ (CRAC) channels by stromal interaction molecule 1 (STIM1). Proc Natl Acad Sci U S A 108, 13299–13304

10. Calloway, N., Owens, T., Corwith, K., Rodgers, W., Holowka, D., and Baird, B. (2011) Stimulated association of STIM1 and Orai1 is regulated by the balance of PtdIns(4,5)P<sub>2</sub>between distinct membrane pools. J Cell Sci 124, 2602–2610

11. West, S. J., Kodakandla, G., Wang, Q., Tewari, R., Zhu, M. X., Boehning, D., and Akimzhanov, A. M. (2022) S-acylation of Orai1 regulates store-operated Ca2+ entry. J Cell Sci 135

12. Carreras-Sureda, A., Abrami, L., Ji-Hee, K., Wang, W. A., Henry, C., Frieden, M., Didier, M., van der Goot, F. G., and Demaurex, N. (2021) S-acylation by ZDHHC20 targets ORAI1 channels to lipid rafts for efficient Ca2+ signaling by Jurkat T cell receptors at the immune synapse. Elife 10

13. Katz, Z. B., Zhang, C., Quintana, A., Lillemeier, B. F., and Hogan, P. G. (2019) Septins organize endoplasmic reticulum-plasma membrane junctions for STIM1-ORAI1 calcium signalling. Sci Rep 9, 10839

14. Qin, X., Liu, L., Lee, S. K., Alsina, A., Liu, T., Wu, C., Park, H., Yu, C., Kim, H., Chu, J., Triller, A., Tang, B. Z., Hyeon, C., Park, C. Y., and Park, H. (2020) Increased Confinement and Polydispersity of STIM1 and Orai1 after Ca(2+) Store Depletion. Biophys J 118, 70–84

15. Mitchell, D. A., Vasudevan, A., Linder, M. E., and Deschenes, R. J. (2006) Protein palmitoylation by a family of DHHC protein S-acyltransferases. J Lipid Res 47, 1118–1127

16. Ohno, Y., Kihara, A., Sano, T., and Igarashi, Y. (2006) Intracellular localization and tissue-specific distribution of human and yeast DHHC cysteine-rich domain-containing proteins. Biochim Biophys Acta 1761, 474–483

17. Aicart-Ramos, C., Valero, R. A., and Rodriguez-Crespo, I. (2011) Protein palmitoylation and subcellular trafficking. Biochim Biophys Acta 1808, 2981–2994

18. Chen, J. J., Fan, Y., and Boehning, D. (2021) Regulation of Dynamic Protein S-Acylation. Front Mol Biosci 8, 656440

19. Emrich, S. M., Yoast, R. E., Xin, P., Arige, V., Wagner, L. E., Hempel, N., Gill, D. L., Sneyd, J., Yule, D. I., and Trebak, M. (2021) Omnitemporal choreographies of all five STIM/Orai and IP3Rs underlie the complexity of mammalian Ca(2+) signaling. Cell Rep 34, 108760

20. Dynes, J. L., Amcheslavsky, A., and Cahalan, M. D. (2016) Genetically targeted single-channel optical recording reveals multiple Orai1 gating states and oscillations in calcium influx. Proc Natl Acad Sci U S A 113, 440–445

21. Feske, S. (2007) Calcium signalling in lymphocyte activation and disease. Nat Rev Immunol 7, 690–702

22. Shaw, P. J., and Feske, S. (2012) Regulation of lymphocyte function by ORAI and STIM proteins in infection and autoimmunity. J Physiol 590, 4157–4167

23. Pacheco, J., Dominguez, L., Bohórquez-Hernández, A., Asanov, A., and Vaca, L. (2016) A cholesterol-binding domain in STIM1 modulates STIM1-Orai1 physical and functional interactions. Sci Rep 6, 29634

24. Zheng, S., Zhou, L., Ma, G., Zhang, T., Liu, J., Li, J., Nguyen, N. T., Zhang, X., Li, W., Nwokonko, R., Zhou, Y., Zhao, F., Liu, J., Huang, Y., Gill, D. L., and Wang, Y. (2018) Calcium store refilling and STIM activation in STIM-and Orai-deficient cell lines. Pflugers Arch 470, 1555–1567

25. Bieerkehazhi, S., Fan, Y., West, S. J., Tewari, R., Ko, J., Mills, T., Boehning, D., and Akimzhanov, A. M. (2022) Ca2+-dependent protein acyltransferase DHHC21 controls activation of CD4+ T cells. J Cell Sci 135

26. Tewari, R., Shayahati, B., Fan, Y., and Akimzhanov, A. M. (2021) T cell receptor-dependent S-acylation of ZAP-70 controls activation of T cells. J Biol Chem 296, 100311

27. Fan, Y., Shayahati, B., Tewari, R., Boehning, D., and Akimzhanov, A. M. (2020) Regulation of T cell receptor signaling by protein acyltransferase DHHC21. Mol Biol Rep 47, 6471–6478

28. Akimzhanov, A. M., and Boehning, D. (2015) Rapid and transient palmitoylation of the tyrosine kinase Lck mediates Fas signaling. Proc Natl Acad Sci U S A 112, 11876–11880

29. Pani, B., and Singh, B. B. (2009) Lipid rafts/caveolae as microdomains of calcium signaling. Cell Calcium 45, 625–633

30. Quintana, A., Pasche, M., Junker, C., Al-Ansary, D., Rieger, H., Kummerow, C., Nunez, L., Villalobos, C., Meraner, P., Becherer, U., Rettig, J., Niemeyer, B. A., and Hoth, M. (2011) Calcium microdomains at the immunological synapse: how ORAI channels, mitochondria and calcium pumps generate local calcium signals for efficient T-cell activation. EMBO J 30, 3895–3912

31. Yokosuka, T., and Saito, T. (2010) The immunological synapse, TCR microclusters, and T cell activation. Curr Top Microbiol Immunol 340, 81–107

32. Akimzhanov, A. M., Wang, X., Sun, J., and Boehning, D. (2010) T-cell receptor complex is essential for Fas signal transduction. Proc Natl Acad Sci U S A 107, 15105–15110

33. Garcia, M. I., Chen, J. J., and Boehning, D. (2017) Genetically encoded calcium indicators for studying long-term calcium dynamics during apoptosis. Cell Calcium 61, 44–49

34. Tewari, R., West, S. J., Shayahati, B., and Akimzhanov, A. M. (2020) Detection of Protein S-Acylation using Acyl-Resin Assisted Capture. J Vis Exp

